# Fast cortical dynamics encode tactile grating orientation during active touch

**DOI:** 10.1101/2020.07.28.225565

**Authors:** Evan R. Harrell, Anthony Renard, Brice Bathellier

**Affiliations:** Department for Integrative and Computational Neuroscience (ICN) Paris-Saclay Institute of Neuroscience (NeuroPSI) UMR9197 CNRS/University Paris Sud CNRS, Building 32/33, 1 avenue de la Terrasse, 91190 Gif-sur-Yvette, France; Institut Pasteur, INSERM, Institut de l’Audition, 63 rue de Charenton, F-75012 Paris, France

## Abstract

Touch-based object recognition relies on perception of compositional tactile features like roughness, shape, and surface orientation. However, besides roughness, it remains unclear how these different tactile features are encoded to guide perception. Here, we establish a barrel cortex-dependent perceptual task in which mice discriminate tactile gratings based on orientation using only their whiskers. Multi-electrode recordings in barrel cortex reveal weak orientation tuning in average firing rates during grating exploration despite high levels of cortical activity. Just before decision, orientation information extracted from fast cortical dynamics more closely resembles concurrent psychophysical measurements than single neuron orientation tuning curves. This temporal code conveys both stimulus and choice-related information, suggesting that fast cortical dynamics during exploration of a tactile object both reflect the physical stimulus and impact the upcoming perceptual decision of the animal.

## Introduction

Touch-based object recognition is essential for guiding behavior in a wide variety of environmental conditions. Reliable recognition generally depends on tactile search behavior executed with appendages like fingers for humans or the mystacial vibrissae for rodents^1^. The vibrissae, or whiskers, are rooted on the rodent snout in densely innervated follicles, where mechano-sensitive cells transduce whisker bending and contact forces into electrical signals^2^. The resulting sensory information has spatial (across whiskers) and temporal aspects that are integrated as it passes through several distinct somatosensory pathways before reaching barrel cortex and other areas^3^. As the foremost recipient of primary somatosensory thalamic afferents^4^, barrel cortex is seen as the major cortical hub for the processing of whisker-based tactile information^5,6^. However, its precise functional roles remain poorly understood, as it has been difficult to disentangle the multiplexed encoding of whisker touches and self-generated movement^7,8^.

Extensive studies on how barrel cortex neurons respond to simple, reliably targeted whisker stimuli have pointed towards a somato-topographical code based on high velocity deflections of single whiskers^9^. However, behavioral studies suggest this simple coding framework is not sufficient to support some of the perceptual functions of barrel cortex. In head-fixed mice, barrel cortex indispensably encodes the precise location of an object in a task requiring whisker search behavior^10,11^. This simple feature, location, is already beyond what a code based purely on velocity can represent for a single whisker. Although barrel cortex is essential to precisely locate objects with the whiskers, it is not required to actively detect the simple presence or absence of objects in the proximal surroundings^12,13^. This detection process can likely be supported by lower brain areas. In more demanding task conditions like the discrimination of sandpapers^14–21^ or in situations that require cognitive planning like whisker-mediated gap crossing^12^, barrel cortex is once again essential. Taken together, perceptual studies suggest that barrel cortex is critical for precisely placing and recognizing tactile objects, especially in conditions that demand spatial and temporal integration of tactile inputs. The simple coding schemes generated from reliably targeted whisker stimuli do not shed light on how barrel cortex serves these perceptual processes.

To better understand the neural underpinnings of tactile object recognition, it is thus crucial to study the temporal evolution of the barrel cortex encoding for a multitude of tactile features during perceptual behaviors. In this pursuit, many studies have focused on how the coarseness of anisotropic surface textures (sandpapers) is encoded during exploration with one or a few whiskers^14–21^. These studies found that object coarseness is encoded by temporal integration of whisker slip events, with higher rates of slip events causing higher firing rates in barrel cortex^14–23^. While this has provided insights into tactile coding, coarseness is just one of many features that can differ between objects. Along with variations in coarseness, natural objects also exhibit unique combinations of isotropic features, which means they can be decomposed into an arrangement of oriented surfaces. While freely moving rats can discriminate oriented tactile gratings with their whiskers^24^, it is not known if and on what time scale grating orientation is encoded in barrel cortex during active sensation. To study this, we developed a barrel cortex-dependent Go/NoGo task in which head-fixed mice discriminate tactile gratings based on orientation with their whiskers. Multi-electrode recordings during task performance revealed that while peak cortical firing rates occurred early on during grating exploration, stable orientation selectivity (in the average firing rates) of individual units in this period was weak. However, orientation-specific temporal sequences of activity could be observed in some neuronal ensembles and support vector machine (SVM) classifiers based on the time course of population activity during grating exploration decoded the orientation category in line with concurrent psychophysical measurements. These results suggest that orientation information is first encoded in the temporal dynamics of population activity in barrel cortex, and only later do individual units develop orientation tuning in their average firing rates. Fast cortical dynamics contain both sensory and choice-related information, as revealed by analysis of the perceptual errors committed by the animals. Thus, the fast cortical dynamics during exploration provide the initial sensory encoding, and their discriminability is then linked with the perceptual choice.

## Results

### Mice categorize texture gratings based on orientation

To investigate if mice are able to discriminate tactile gratings, we trained head-fixed, water-deprived mice (**Fig. 1A**, fig. S1) to report the perceived orientation of a tactile grating by licking a tube to receive a water reward. The oriented gratings were presented in full dark conditions using a linear stage, and whisker interactions with the gratings were filmed with a high-speed infrared video camera (see Methods). For each trial, after no licking was detected on the reward port for at least 3 seconds, a 2 kHz sound was played to signify trial onset and a grating was translated into reach of the right whisker field (**Fig. 1B**). After a 1 second period of interaction with the grating, mice reported the orientation of the grating by either licking to obtain a water reward (Go trial) or refraining from licking to avoid punishment (**Fig. 1B**). In these trial conditions, mice were trained to perform a simple Go/NoGo discrimination between a vertically oriented grating (90°) and a horizontal grating (0°), with Go and NoGo stimulus types interchanged in different groups of animals (fig. S1 and Methods for all training details). After performance of simple Go/NoGo discrimination stabilized above 70% correct across 2 days, intermediate orientation angles spaced by 9° were gradually introduced and reinforced (**Fig. 1B**, see Methods). In this psychometric version of the task, the boundary between rewarded and non-rewarded orientations was 45°, and the fully ambiguous orientation was never presented.

**Figure 1.**
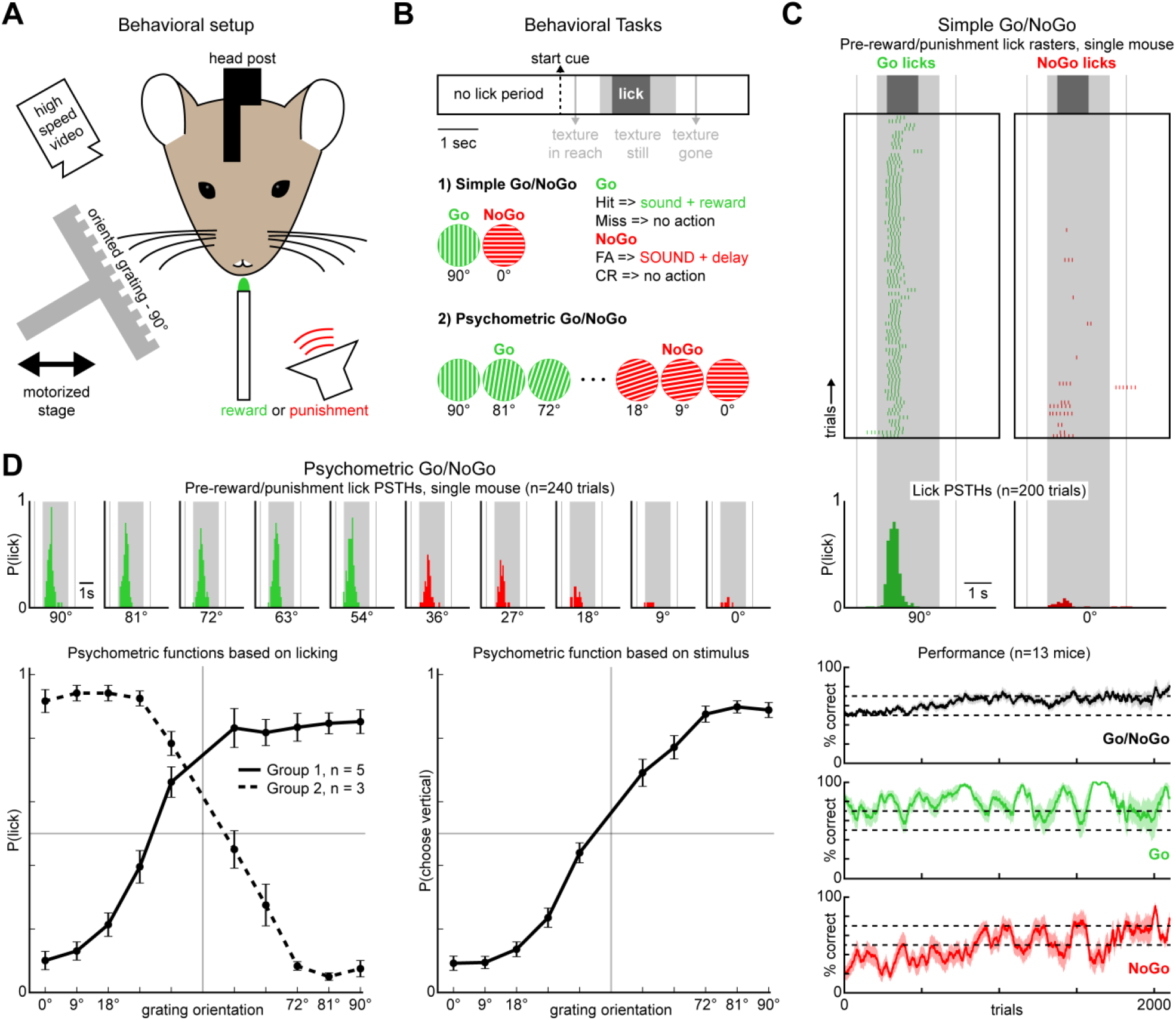
Mice categorize tactile gratings based on their orientation. **A.** A schematic showing the behavioral setup. **B.** Task parameters for (1) simple Go/NoGo discrimination and (2) psychometric Go/NoGo grating orientation discrimination tasks. **C.** Top: Lick raster from simple Go/NoGo discrimination between vertical (90°) and horizontal (0°) gratings. Middle: Lick probabilities from one session of simple Go/NoGo discrimination. Bottom: Mean learning curves (shaded areas are s.e.m., n=13 animals) for All (black), Go (green), and NoGo trials (red). **D.** Top: Lick probabilities from one session of psychometric Go/NoGo discrimination. Bottom left: Psychometric functions for two groups of mice where the Go/NoGo rules were interchanged. Bottom right: The psychometric functions controlled for motivation (error bars are s.e.m.).

From the beginning of simple Go/NoGo (0° vs. 90°) training, mice quickly learned the appropriate time to lick and after 10-15 days (~2000 trials), they discriminated between orthogonal grating orientations as measured by their licking behavior (**Fig. 1C**). Improved performance across time was mostly attributable to refraining from licking for the NoGo stimuli (**Fig. 1C**). After progressing to the psychometric version of the task, the ongoing motivational state of the animal, driven by thirst, determined whether False Alarm or Miss errors were more common. In most animals, we observed a more gradual change in licking behavior across orientation steps for NoGo than for Go orientations **(Fig. 1D**, lick histograms). Along with this, mice tended to make more False Alarm errors than Miss errors (**Fig. 1D**) indicative of a strategy aiming to minimize reward loss. This strategy results in asymmetric psychometric functions (**Fig. 1D**). To balance these curves^25^, we averaged across animals in which the Go and NoGo orientations had been interchanged, and this revealed that the discrimination performance controlled for motivation is almost perfectly symmetric (**Fig. 1D**). These results indicate that head-fixed mice can discriminate tactile gratings using only their whiskers, and they do so with high acuity.

### Barrel cortex is essential for discriminating oriented tactile gratings

We next asked if barrel cortex is essential to perform the simple Go/NoGo version of this task. Optogenetic manipulations can perturb performance even if a brain area is dispensable^13^, so we opted for a barrel cortex lesioning strategy. Mice were trained in the simple Go/NoGo version of the task until they reached stable performance above 70% correct across 2 days, after which thermo-coagulation lesions^26^ were applied over the entire contralateral postero-medial barrel field (**Fig. 2A**, fig. S2). As a control, another group of animals (sham group) underwent mock surgeries that involved the same duration of anesthesia, a large craniotomy over barrel cortex, and the same process to reseal the exposed brain but with no lesion. The day after surgery, both lesion and sham groups performed the simple Go/NoGo task at chance levels (**Fig. 2B**), indicating that the general aftereffects of surgery and craniotomy have an impact on performance. Over the ensuing days, the sham group steadily recovered performance, while the lesioned group continued to perform at chance levels (**Fig. 2B**). Lesions were examined post hoc in coronal sections to assure that all postero-medial barrels (straddlers, A1-E4) in the whisker region of the primary somatosensory cortex had been removed (fig. S2).

**Figure 2.**
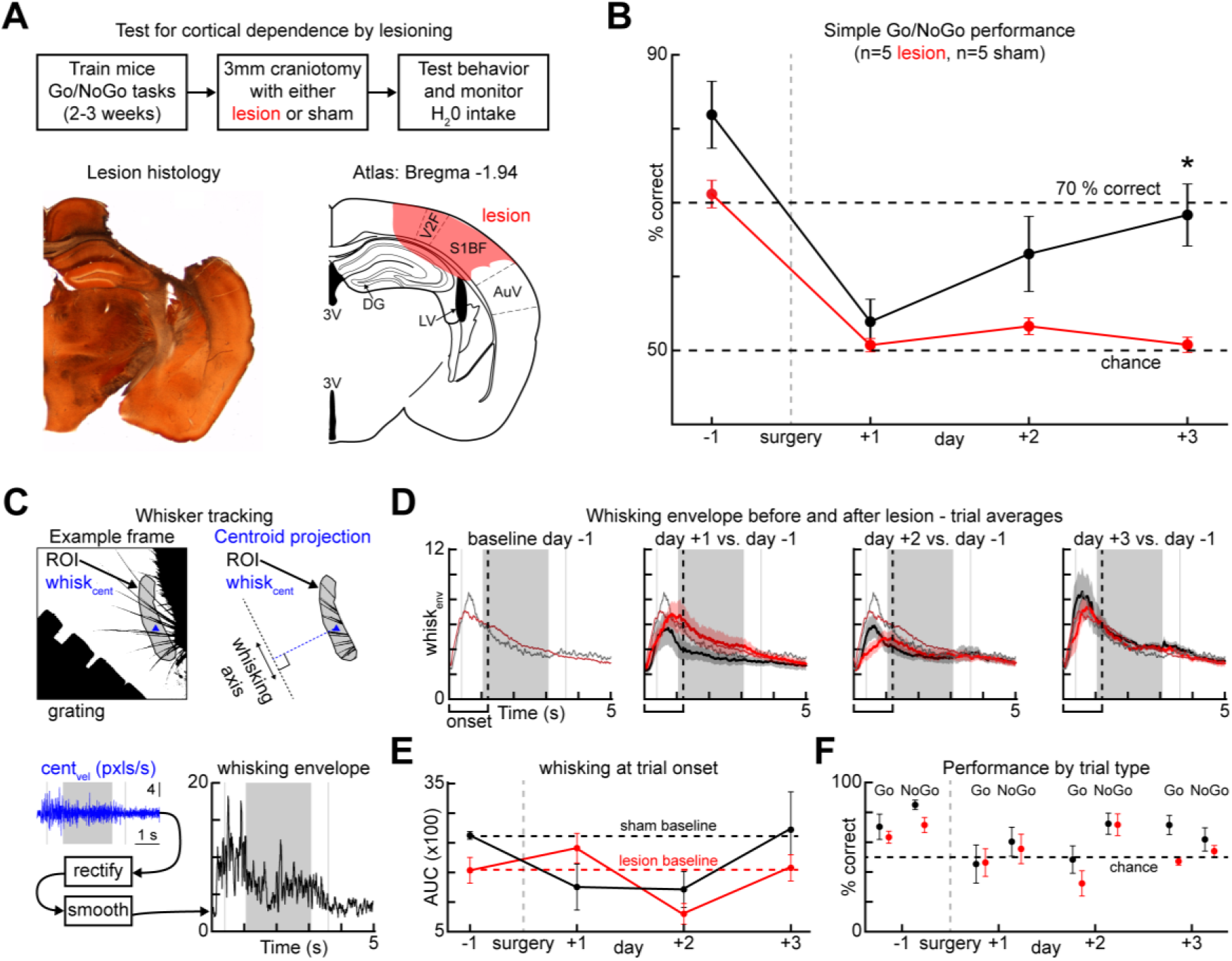
Barrel cortex is required for discriminating tactile gratings. **A.** Top: Experimental approach and timeline. Bottom left: An example barrel cortex lesion. Bottom right: the corresponding slice in the brain atlas. Abbreviations: third ventricle (3V), dentate gyrus (DG), lateral ventricle (LV). **B.** Simple Go/NoGo discrimination performance before and after surgery in lesion and sham groups (p=0.0056, bootstrap resample test). **C.** Whisker tracking during performance of the task. Top left: A binarized frame. Top right: A manually selected region of interest (ROI) containing the bases of the whiskers and the centroid (blue). Bottom left: the velocity of the centroid plotted across a trial. Bottom right: the resulting whisking envelope after rectification and smoothing. **D.** Top: Average whisking envelopes across days for lesion and sham groups. **E.** Area under the curve (AUC) during the whisker search period across days for lesion and sham groups. **F.** Performance broken down by trial type for lesion and sham groups.

Barrel cortex lesions are known to affect whisker movement control^13,27^. Therefore, we examined high-speed videos of whisker movements executed by the animals during task performance in sham and lesion groups. To quantify global whisker movements throughout a trial, we defined the whisking envelope as the rectified and smoothed centroid velocity of the binarized whisker image within a manually traced ROI around the whisker bases (**Fig. 2C**, See Methods). This envelope showed that whisking behavior is most pronounced between trial onset (the trial start sound cue) and the time when the grating is fixed and within reach of the whiskers (**Fig. 2D**, gray shaded rectangles). Surgery affected the average whisking envelope in both sham and lesion groups of animals, as quantified by the sum of all whisking at trial onset (**Fig. 2E**, area under the whisking envelope curve). By day 3 after surgery, the total whisking of both groups returned to pre-surgical levels, but the behavioral performance recovered only in the sham group. Therefore, the drop in task performance after lesion cannot be explained by deficiencies in global whisker control. Barrel cortex removal also did not impact performance by abolishing licking. On day 3 after surgery, hit rates and false alarm rates were equal in the lesioned animals (both at ~50%), indicating that mice randomly licked rather than never licking at all (**Fig. 2F**). These results indicate that intact barrel cortex is required to discriminate grating orientations with the whiskers, and this cannot be explained by changes in global whisker search behavior or licking ability.

### Discrimination performance correlates with exploratory whisking and increased barrel cortex spiking activity

To study the encoding of grating orientation in barrel cortex during active discrimination, we made acute extracellular recordings (9 recordings, 74 single unit and 274 multi-units) during the psychometric Go/NoGo version of the task (**Fig. 3A**, fig. S3). Silicon probes with linearly spaced electrodes (spanning 775 μm) were lowered to 1 mm depth from the surface of the contralateral barrel cortex (targeted C2 whisker A/P: −1.5mm, M/L: 0/3.3mm). Electrode placement in the barrel cortex was histologically verified in tangential sections after the experiments (fig. S3), and most of the active cells that were recorded resided in deeper layers (fig. S3). All recorded mice showed stable task performance above 70% correct on the day before the recording, but only some of them went on to perform during the recording (n=5 discriminating mice), while others did not (n=4 non-discriminating mice). This is likely due to the anesthesia, durotomy, and electrode descent preceding behavioral measurements in these acutely recorded animals.

**Figure 3.**
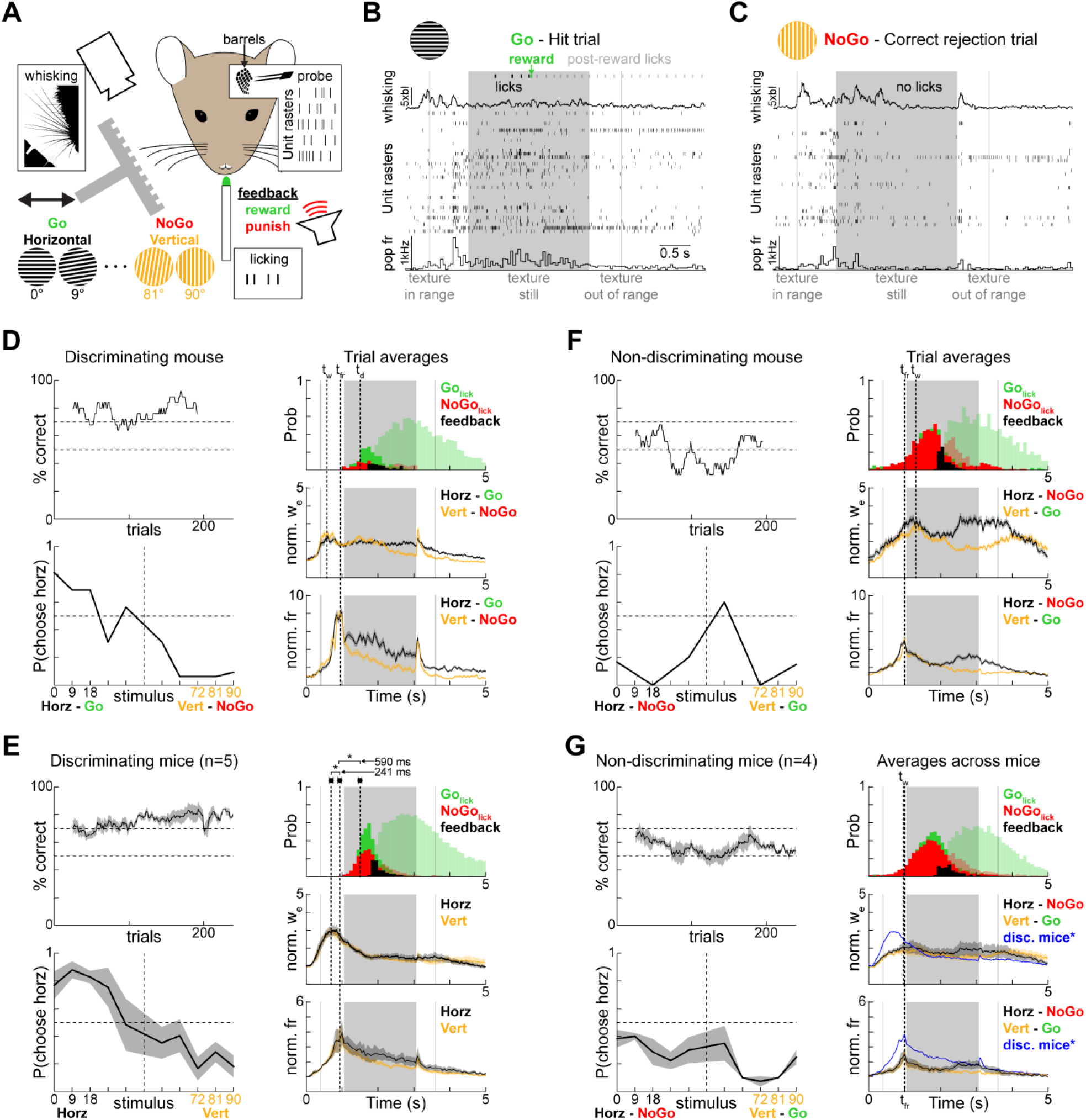
Discrimination performance correlates with exploratory whisking and increased barrel cortex spiking activity. **A.** A schematic of the recording setup. **B.** An example hit trial showing licks, reward, whisking, and unit rasters. **C.** Same as B for a correct rejection trial. **D-E.** Performance across trials, psychometric functions, licking, whisking, and total population spiking activity for a discriminating mouse (D) or 5 discriminating mice (E). Shaded lick histograms are licks after reward/punishment. Shading around curves is s.e.m. t_w_ is the time that the trial averages of whisking were at a peak, t_fr_ is the peak firing rate, and t_d_ is the time that licking became discriminative. **F-G.** Same as D-E but for non-discriminating mice.

In an example hit trial from a discriminating animal (**Fig. 3B**), the mouse initiated whisking before the grating came into reach and spiking activity increased once the grating was close enough to touch the whiskers. After ~500 ms of exploration, the mouse decided to lick and received a water reward, which triggered prolonged licking. In an example correct rejection trial (**Fig. 3C**), the mouse whisked into the grating, which produced spiking activity in the last ~250 milliseconds before the grating stabilized at its fixed position in reach. Then, the mouse correctly withheld licking to avoid punishment. This same behavioral sequence was apparent when averaging across all trials in this animal (**Fig. 3D**) or across all discriminating animals (**Fig. 3E**). As the grating approached the mice, they executed whisker search behavior, which was followed by a burst of spiking activity in barrel cortex neurons that peaked just before the grating stopped near the snout. Licking behavior was initiated after the grating stopped and the average licking differences between orientation classes (> or < 45°) could be discriminated ~590 ms after the peak of population activity (**Fig. 3E**, see Methods). After the decision to lick, low whisking levels were maintained and, in some mice, a rebound of whisking and barrel cortex activity was observed when the texture retraction was initiated (**Fig. 3E**).

These patterns of behavior were much less discernible in animals that did not discriminate the gratings during the recording (**Fig. 3F-G**). In these animals, licking was initiated earlier, even before the grating came to a halt, suggesting that their choice behavior did not take the grating into account. Whisking levels and spiking activity were also reduced, especially during the early exploration of the grating. However, the population firing rates still peaked just before the grating arrived at its fixed position, suggesting that there are still sensory responses even if these animals did not perform the discrimination. These data indicate that patterned behavior (search, find, lick) and increased spiking activity in barrel cortex were associated with task performance.

### Orientation tuning in average firing rates is weak during grating exploration

To study how barrel cortex encoded grating orientation, we first examined individual unit activity. Some single units discharged many spikes at the onset of whisker interactions with the grating for all different orientations, and their firing rates decreased when the grating reached its fixed position (**Fig. 4A**, fig. S3, Single Unit 1). Other neurons had less pronounced responses during early exploration, but still had elevated firing rates while the gratings were within reach (**Fig. 4A**, fig. S3, Single Unit 2). To quantify whether unit responses were selective for a particular grating orientation, we constructed tuning curves in 500 ms windows at different latencies with respect to trial onset (**Fig. 4B**, fig. S4, blue-magenta gradient). Tuning curves were computed for units by summing the spikes within a 500 ms window of interest for each trial of a given orientation, then dividing the total number of spikes for each trial by the size of the window (**Fig. 4B**, fig. S4, 6 examples). To assess tuning significance, we expressed the tuning curves in polar form and compared the magnitude of the vector sum to the vector sums obtained from shuffling the trial labels 200 times (**Fig. 4B** bottom left). If the actual tuning vector was beyond the 95^th^ percentile of the shuffled distribution, the unit was considered significantly tuned. For example, single Unit 1 showed no selectivity during the first interactions with the grating (**Fig. 4B** top left, blue) or after discriminative choice (**Fig. 4B** top left, magenta). Single Unit 2 was tuned to grating orientation (**Fig. 4B** right). The tuning to 18° gratings originated in the second 500 ms time bin (before decisive licking in this animal measured across all trials, see Methods) after the grating came into reach of the whiskers, and it persisted through decision and feedback.

**Figure 4.**
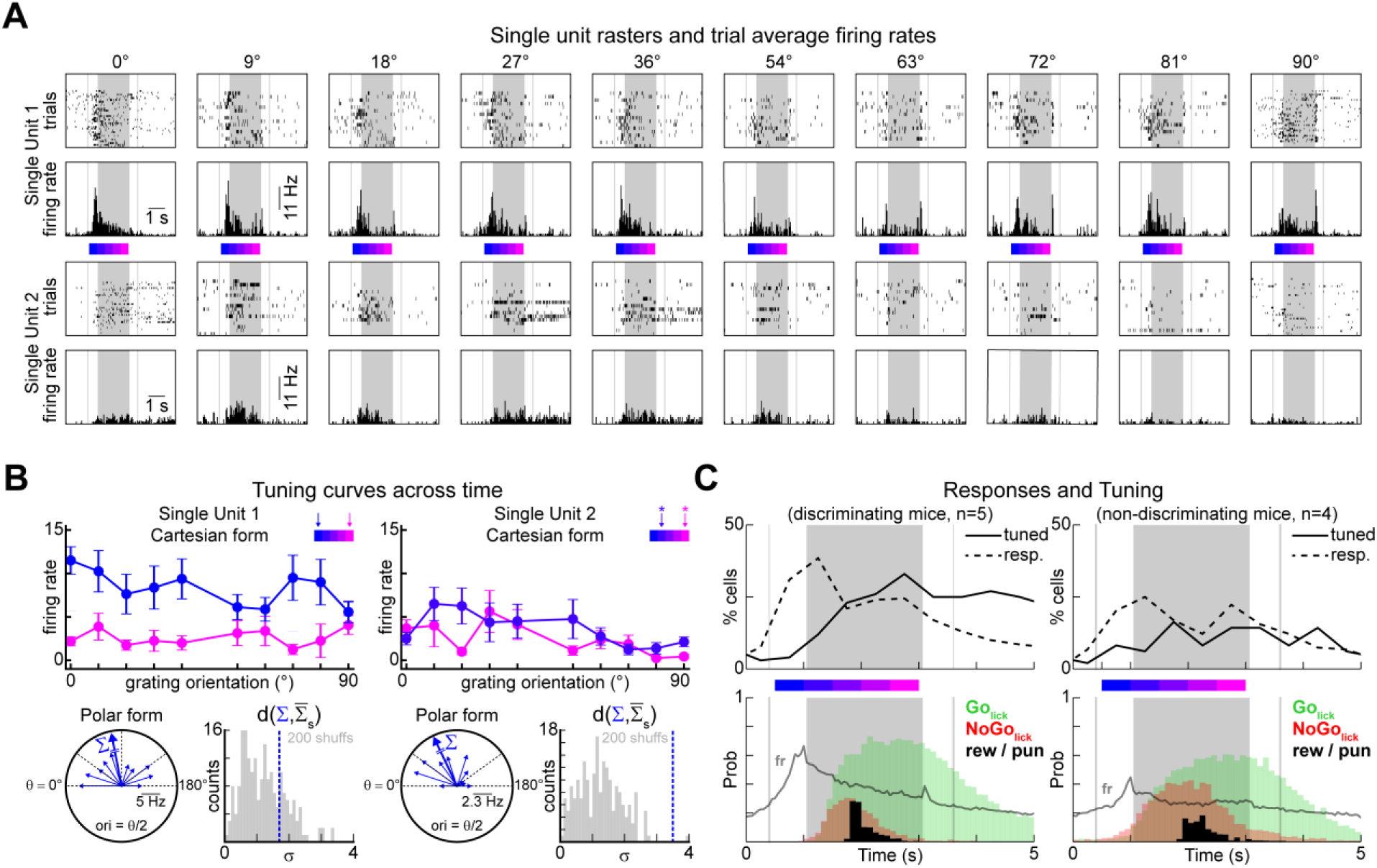
Orientation tuning is scarce during early grating exploration. **A.** Single trial rasters and trial-averaged PSTHs from two example single units during performance of the psychometric Go-NoGo task. **B.** Tuning curves from 2 example single units (same units from a) in various time windows color-coded by where the 500 ms time bin in which the curves were computed falls with respect to early vs. late periods. **C.** The percentage of responsive cells and orientation-tuned cells relative to trial onset. Population firing rates and licking behavior are plotted below to serve as a reference.

Examining the orientation tuning across time for all units, we found that it was not above chance levels during the peak of firing that occurs during the first interactions with the grating (just before grating halt), and faintly started to appear in the period between grating halt and discriminative licking (**Fig. 4C**). This suggests that in the exploratory period, information about grating orientation is not yet well-encoded by average firing rates of single neurons, as might be the case in other sensory modalities^28^. In non-discriminating animals, not only were there fewer responsive cells in barrel cortex, but also fewer tuned cells in all periods (**Fig. 4C** bottom right). Taken together, these results suggest that orientation selectivity in the firing rates of individual units is scarce before decision.

### Temporal decoders outperform rate decoders during grating exploration and provide the closest match to psychophysical measurements

Because tactile inputs occur in a sequence of multi-whisker contacts that evoke dynamic cortical responses, we hypothesized that during exploration before decision, grating orientation information was carried by coordinated population activity sequences rather than by the firing rate of orientation-tuned units. To search for orientation-specific activity sequences, we looked at the co-firing patterns for all simultaneously recorded units in an example mouse. Co-firing matrices were computed separately for horizontal and vertical trials in the time period leading up to grating halt (**Fig. 5A**, top, 5×100ms bins). Out of the 37 simultaneously recorded units in this example mouse, 9 had stronger co-firing for horizontal gratings and 8 neurons had stronger co-firing for vertical gratings in the chosen time period (**Fig. 5A**, top, red and blue regions in the co-firing matrix), while the rest of the population had equal co-firing for horizontal and vertical trials. We examined the time course of the trial-averaged response separately for these two subgroups of neurons, and we observed distinct temporal sequences of activity for vertical and horizontal trials in both sub-populations (**Fig. 5A**, bottom, strongest in the group co-activated by horizontal gratings). This observation suggests that co-activation sequences might carry some information about grating orientation.

**Figure 5.**
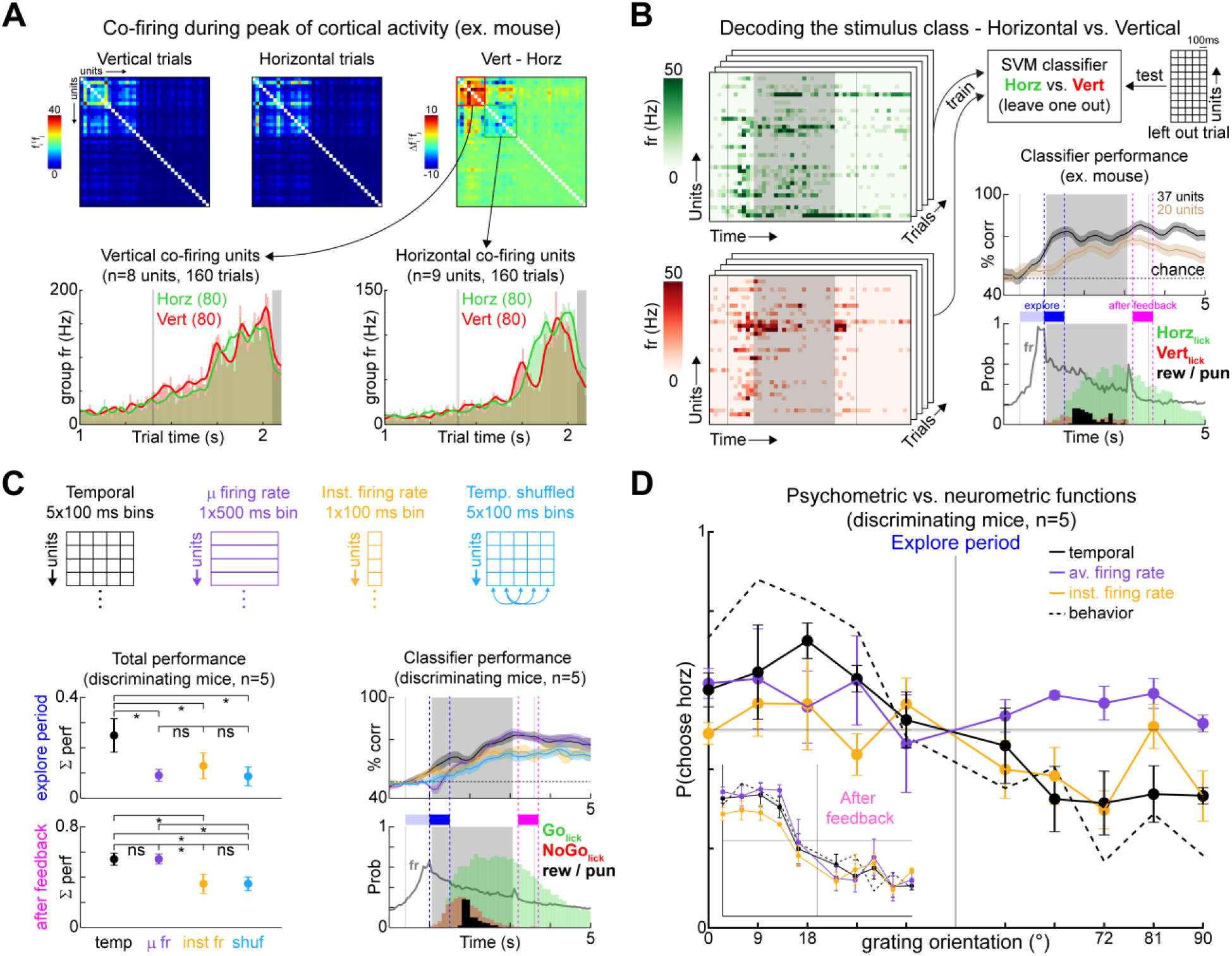
Temporal decoders outperform rate decoders during grating exploration and provide the closest match to psychophysical measurements. **A.** Top: Co-firing matrices from an example population (n=37 units) as the grating approached. After computing the difference between vertical and horizontal matrices, two groups of units were apparent. Bottom: Time course of group firing rate as grating approached. **B.** Schematic showing how the support vector machine (SVM) classifiers were trained and tested for one example mouse. Classifier performance is aligned with the licking behavior and the population firing rates. The end of the exploratory period is defined as the 500 ms before discriminative licking (across all trials) and the after-feedback period is the same size time window after reward or punishment. **C.** Four classifier types defined by their bin arrangements were tested to assess the contribution of fast cortical dynamics in the sample populations of neurons. Performance, licking behavior and population firing rates are all aligned. Total performance is the area under the curve that is above chance in a period. * means p<0.05 in a bootstrap resample test. **D.** Psychometric and neurometric functions for the three main classifier types in the exploratory period or after feedback (inset).

To generally examine whether temporal patterns of population activity contain grating information, we trained support vector machine (SVM) classifiers which take as input the number of spikes that each unit in the population produced in the 5 preceding 100ms time bins (**Fig. 5B**, same example mouse). After leave-one-out cross-validation, the performance of this temporal decoder in classifying vertical versus horizontal gratings was above chance well-before the onset of licking or orientation tuning (**Fig. 4**). Classifier performance before decision was drastically reduced by removing the 17 units with orientation-specific co-firing patterns (**Fig. 5B**, brown), and training only on these 17 units gave performance that was indistinguishable from the whole population (fig. S5), confirming that the observed co-firing patterns provide orientation information during early grating exploration. To broadly evaluate the importance of temporal patterns of activity, we compared the ability of 4 different decoders to discriminate horizontal and vertical gratings in all task-performing mice: (i) the temporal decoder using the sequence of activity over the preceding 5×100ms time bins, (ii) a mean firing rate decoder using the sum of activity over a single preceding 500ms time bin, (iii) an instantaneous firing rate decoder taking activity over a single preceding 100ms time bin, and (iv) a decoder using the same 5×100ms bin sequence as the temporal decoder but with the time bins shuffled across trials to control for dimensionality but destroy temporal patterning. Across all mice, the decoders with fine temporal resolution outperformed the other decoders during exploration, with the best performance coming from the temporal decoder with a 5-bin history. Increasing the temporal resolution of the classifier binning did not affect these conclusions (fig. S5). This demonstrates that temporal sequences of population activity contain more information about grating orientation than the average firing rates before decision. This advantage vanished after decision and feedback, as the appearance of orientation tuning and differences in licking behavior drove discernible differences in cortical activity (**Fig. 5C**, **Fig. 4**).

We next examined which decoder best followed concurrent psychophysical measurements by looking at classifier performance across different grating orientations (**Fig. 5D**). During exploration, the temporal decoder again out-performed rate decoders and showed the closest resemblance to the psychometric behavior (**Fig. 5D**, right). This indicates that precise temporal co-activation patterns provide a sensory coding space that can underlie this perceptual behavior. After decision, temporal and rate decoders were equally predictive of psychophysical measurements, which could either reflect the pronounced differences in licking behavior between grating classes in this time window or a reformatting of the sensory code.

### Temporal sequences of cortical activity during grating exploration encode sensory as well as choice-related information

We next examined if the activity preceding decision in barrel cortex reflects the external sensory object or the perceptual decision of the animal by measuring to what extent it predicts single trial outcomes, which in our case were Hits, Misses, False Alarms and Correct Rejections (**Fig. 6A**). Because most animals performed very few Misses, we concentrated on Hit, False Alarm, and Correct Rejection trials. When averaging across all discriminating animals, there were no differences in whisking behavior and population spiking activity during grating exploration for these different trial outcomes, however there was more whisking and increased barrel cortex activity after punishment for False Alarms (**Fig. 6B**).

**Figure 6.**
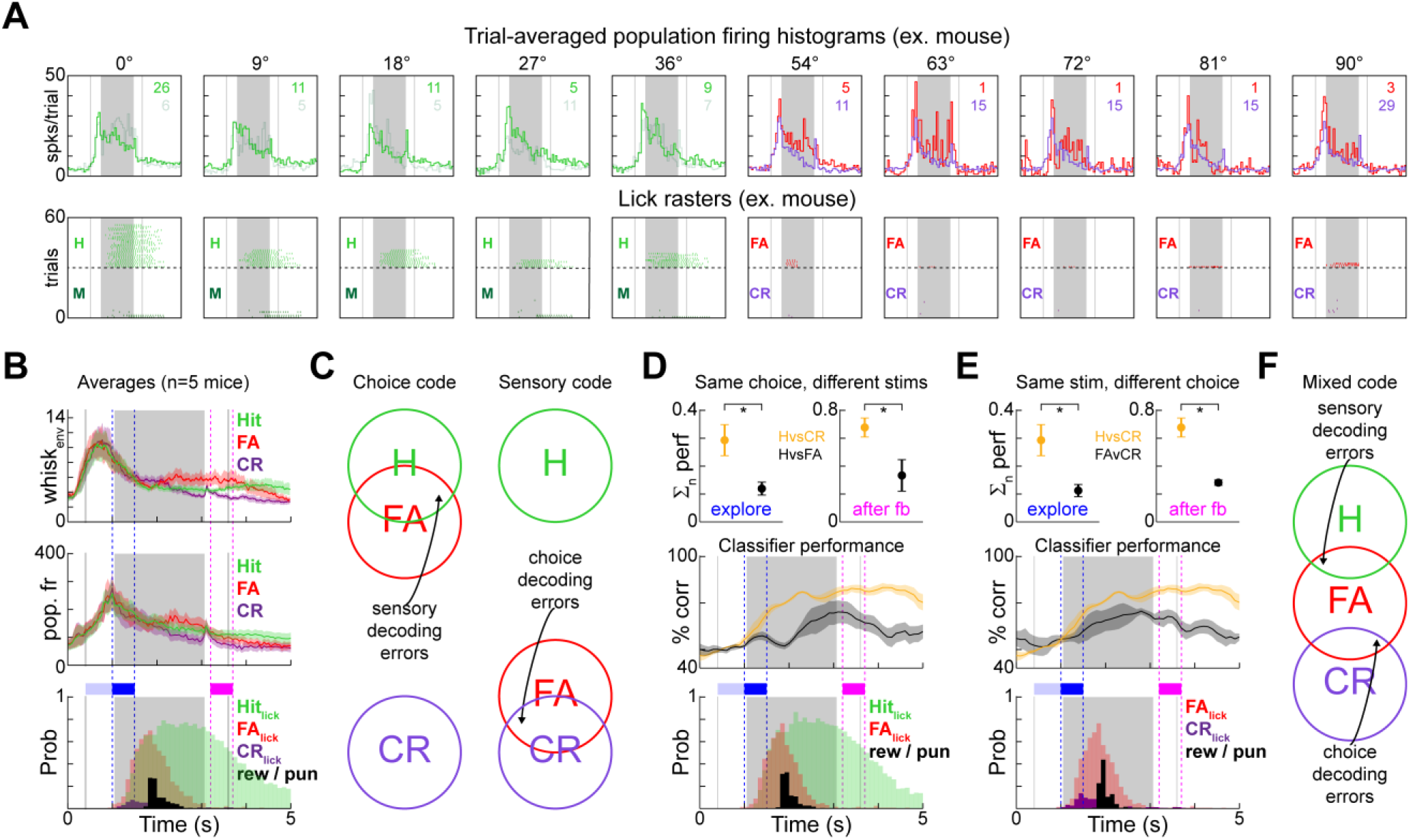
Temporal sequences of cortical activity during grating exploration encode sensory as well as choice-related information. **A.** Top: Trial-averaged population firing rates separated by orientation and outcome for an example mouse (Hits, Misses, False Alarms, and Correct Rejections are lime green, dark green, red, and purple respectively) Bottom: Lick rasters for all trials for the same mouse. **B.** Left: Trial-averaged whisking envelope, population firing rate, and licking behavior separated for Hits (lime green), Correct Rejections (purple), and False Alarms (red) for all discriminating mice. **C.** Schematics for a choice code and a sensory code. **D.** SVM classifier decoding for Hits vs. False Alarms (black) and Hits vs. Correct Rejections (gold). Top: total performance in the exploratory and after feedback periods for the different classifiers normalized to the baseline before trial start (p=0.0214 and p=0.035, paired bootstrap resample test), Middle: Classifier performance across all time points relative to trial onset. Bottom: Licking behavior for Hits and False Alarms. **E.** Same as D for False Alarms vs. Correct Rejections (p=0.018 and p=0.0134, paired bootstrap resample test). **F.** Schematic for a mixed code, which is observed to be the case in S1.

If barrel cortex encoded only the choice and not the sensory object, which could diverge due to various encoding or decision error sources, then Hit and False Alarm trials should be indistinguishable in the cortical activity during exploration (**Fig. 6C**). On the contrary, if barrel cortex faithfully encoded the sensory object regardless of the choice, then Correct Rejection and False Alarm trials should be indistinguishable in the cortical activity during the same period (**Fig. 6C**). When testing these scenarios, we found that temporal decoders trained to discriminate Hit vs. False Alarm trials performed worse in the exploratory period than decoders trained to discriminate Hit vs. Correct Rejection trials (**Fig. 6D**). This suggests that the False Alarm errors were at least partly due to sensory encoding errors in which the ‘No-Go’ sensory object was misrepresented as being more like a ‘Go’ sensory object, prompting the animal to lick. But classifier performance for Hit vs. False Alarm trials in the exploratory period was not completely abolished, suggesting that there is another component that reflects the sensory object regardless of choice. In line with this, temporal decoders performed worse at discriminating False Alarm vs. Correction Rejection trials for matched stimuli than they did at discriminating Hit vs. Correct Rejection trials (**Fig. 6E**). This confirms that sensory information is also present in barrel cortex that does not necessarily lead to correct choices, and also rules out the possibility that licking differences alone can drive classifier performance. Taken together, these analyses establish that temporal sequences of population activity in barrel cortex encode both sensory and choice-related information during grating exploration (**Fig. 6F**).

## Discussion

We have shown that head-fixed mice can discriminate tactile gratings based on orientation using only their whiskers (**Fig. 1**), and that this perceptual task is barrel cortex-dependent (**Fig. 2**). Our findings indicate that before a decision has been made, there is more information about grating orientation in the fine temporal dynamics of barrel cortex population activity than in mean firing rates of individual orientation-selective neurons, and decoding this information at higher temporal resolution better reflects the psychophysics of an animal’s perceptual decisions (**Fig. 4**–**6**). This means that the high temporal resolution afforded by electrophysiological recordings is likely to be crucial for understanding the recognition of compound tactile objects composed of sets of oriented edges, because as we have shown, fast temporal variations in barrel cortex activity provide important parts of the initial object representation in S1. This is in stark contrast with the well-studied single whisker detection tasks^13,29,30^, which can easily be solved based on mean firing activity of single barrel cortex neurons. In more challenging task conditions, we propose that tactile information in S1 first appears in fast temporal fluctuations which are further integrated to produce the final decision. There are some precedents to this concept in both rodents and primates. During whisker-based coarseness discrimination, precise spike timing and average firing rates in barrel cortex neurons convey stimulus information that is used by rats to guide decisions^31^. However, the precise form in which the information appears first in barrel cortex was never established, and here we show that the temporal code is indeed present first. In flutter discrimination tasks in macaques, the majority of S1 neurons finely track the precise frequency of the vibration, and thus encode with high temporal resolution^32^. But a relatively smaller number of cells are less locked to the vibration, and instead linearly increase their average firing rates with increasing vibration frequency^32^. Therefore, both temporal and average firing rate codes in S1 can provide the information required for the monkey to solve the task, and psychometric behavioral analysis suggests that the monkeys rely on a mix of the two, because they underperform the ideal observer of the temporal code and outperform the ideal observer of the rate code^32^. Along with these studies, our findings point towards a model where temporal information initially present in S1 is likely transformed either within the primary cortex itself or in concert with other areas to generate integrated firing rate representations, and both of these coding spaces are then linked with tactile perception.

To discriminate tactile gratings, mice need to execute a sequence of appropriate behaviors, and our head-fixed conditions ensured that these behaviors are precisely locked with trial cues. First, mice need to detect the incoming grating. In this pursuit, we found that discriminating mice whisked vigorously when the grating was approaching (**Fig. 3**), and this active search behavior generated temporally dynamic responses in barrel cortex neurons (**Figs. 3**–**5**). Freely behaving rodents can use head movements along with exploratory whisking to perform tactile search^33^, so the level of vigorous whisking observed here might be an adaptation to head-fixed task conditions which could emphasize temporal aspects of the coding. Once the approaching grating is placed, mice continue their search behavior and adaptively sample the grating. We focused our analysis on the time period between first contact and decision, which allowed us to show that orientation tuning in the firing rates of single neurons was scarce (**Fig. 4**) while temporal sequences of population activity could be used in decoding schemes that gave a closer match with concurrent psychophysical measurements. When we examined choice behavior in single trials, we found that the temporally dynamic code in barrel cortex contains both sensory (what the object was) and choice-related (what did the mouse think the object was) information (**Fig. 6**). This suggests that some components of barrel cortex population activity drive the choices made by the animal on a trial by trial basis, as False Alarms tended to occur when cortical activity was less discriminative (**Fig. 6**). This is consistent with detection studies showing that barrel cortex neurons encode both stimulus- and choice-related information^29^. However, our results show that this concept still holds in a more challenging perceptual task in which barrel cortex is essential, which might not be the case for detection^12,13^.

Discriminating oriented gratings with multiple whiskers is a much more challenging task than detecting the vibration or contact of a single whisker, which is reflected by the essential role of the cortex in this task and the temporally dynamic encoding that is present in cortex during task performance. Stimulus complexity aside, when all whiskers can be used, sensory information is spread across the whisker pad. While this lack of control is not ideal for precisely quantifying single whisker contributions to perception, we are still able to recover both sensory and choice-related information from sampled barrel cortex populations. This spread of information across sensors more closely resembles the computational problems that barrel cortex faces in natural conditions than the vibration of a single whisker. The fact that perceptual decisions can still be linked with the discriminability of sampled barrel cortex populations suggests that both sensory and choice-related information are widespread in the cortex even if the same whiskers are not involved on every trial. In summary, our results establish a cortex-dependent tactile discrimination task in which the fast cortical dynamics are informative and lay out how temporally structured co-firing events in subpopulations of neurons can support grating perception. There is much to learn about the circuits that are responsible for transforming temporal population codes into stable firing rate codes^34^ that can be found in downstream areas^24^ and how the interaction between these brain areas facilitates tactile object recognition.

## Materials and Methods

### Animal Care

All experiments were performed in accordance with the French Ethical Committee (Direction Générale de la Recherche et de l’Innovation) and European legislation (2010/63/EU). Procedures were approved by the French Ministry of Education in Research after consultation with the ethical committee #59 (authorization number 9714-2018011108392486). Mice were housed in cages in groups of 2-4 individuals with food available ad libitum on a 12/12 light-dark cycle with temperature kept at 23° C.

### Behavioral Setup

Mice were trained in a custom-built behavioral setup that was interfaced using a National Instruments (NI) card (USB-6343) to control a linear stage (Newmark eTrack series) that brought the gratings within reach of the whiskers and an Arduino Uno to control stepper motors (Makeblock) for adjusting the orientation angle of the grating and a solenoid valve (LVM10R1-6B-1-Q, SMC) for delivering water rewards (5-8 μL). Sound cues were played with loudspeakers (Labtec Spin 85 speakers). Licking signals were acquired and digitized using a capacitive sensor (Sentronic AG, SK-3-18/2,5-B-VA/PTFE) before being fed into the NI card. Software to carry out the training protocols and log the licking data was coded in Matlab using the data acquisition toolbox. Code is available upon request.

### Headpost Implantation

To stabilize the animals in the behavioral apparatus, a head-fixation post was implanted along the mid-line of the skull. Mice (C57BL/6) that were 6-8 weeks old (20-26g) were anesthetized by intraperitoneal injection of a mix of ketamine (Ketasol, 80 mg/kg) and medetomidine (Domitor, 1 mg/kg). Once the mice were insensitive to hindpaw pinch, they were placed on a nose clamp and their eyes were kept moist with Ocrygel (TVM Lab). Body temperature was maintained at 36° using a thermal blanket. Xylocaine was injected under the skin in the center of the skull near bregma. Fur in the surgical location was removed using Veet, and a long incision was made in the skin along the midline of the skull 10 minutes after Xylocaine injection. After being fully exposed, the dorsal surface of the skull was scratched with a scalpel to create striations. The scratched skull was then cleaned with hydrogen peroxide. A head-fixation post was glued in place along the midline using cyanoacrylate, and then the exposed skull and base of the post were covered with Super-Bond (C&B, Sun Medical Co., Ltd.). The implant and all exposed surfaces were then embedded in dental cement. After everything had solidified, the mice were injected in one of the hindlimbs with 15 μL of atipamezole (Antisedan, Orion pharma) and transferred to a recovery cage that was placed on a heating blanket. Mice recovered for at least 1 week before any further manipulation.

### Orientation discrimination training protocol

Mice were weighed every day during water deprivation periods to make sure they did not fall below 80% of pre-deprivation body mass. For two days before training, mice were fully water deprived. On the first day of training, the mice were placed on the head-fixation post for 10 minutes in the dark with the lick port in reach. They were then given single water rewards (5-8 μL) randomly until they started to lick regularly at the lick port. Once they were comfortable licking the lick port for water reward, a protocol was launched that made one reward possible every 10 seconds if the animal licked to initiate the delivery, for up to a maximum of 100 rewards. After this habituation (1-2 days, 1 hour per day), the animals were given trials only with the Go grating until they licked regularly at the correct time within single trials. The trial timeline is shown in **Fig. 1**. For the first 40 trials, rewards were given automatically 1 second after the grating came into reach of the whiskers. After these free rewards, the mice had to lick in a 2 second window that started 1 second after the grating came into reach to receive the reward. The starting threshold to trigger reward was a single lick, which was then increased to as high as 4 (2-4 across all mice) licks to trigger a reward. If animals performed 3 misses in a row, the next Go trial automatically was rewarded, and this ‘miss’ counter was reset while the trial was still scored as a miss. Once the rewards were action-contingent within the trial framework, performance was tracked. When the animals were able to perform 70% correct across an entire training session, a NoGo stimulus was introduced the next day interleaved pseudo-randomly with the Go grating at a ratio of 3 Go trials for every 1 NoGo trial. If the addition of NoGo trials and their associated punishments (white noise at 60-70 dB and time out of 5-20 seconds) did not cease reward seeking behavior, the ratios were equilibrated (50% Go 50% NoGo) on the next day of training. The first NoGo stimulus was a flat surface (a small circle of printer paper glued on a disk the same dimensions as the gratings) with no grating (fig. S1). Once the animals discriminated this flat surface from the Go grating (fig. S1, performed 70% correct across 200 trials in a single day), the NoGo stimulus was changed to a grating orthogonal to the Go grating. Punishments (loudness of the white noise and length of the time out) and lick thresholds were increased if animals could not refrain from licking for NoGo gratings. After 2 days of 70% performance in discriminating orthogonal gratings, intermediate grating orientations were introduced. At first, only 4 intermediate orientations (9, 18, 72, and 81°) were given, but then another 4 (27, 36, 54, and 63°) were added after performance stabilized above 70% correct. For the full psychometry, a single training session contained 40 trials for each extremity (0 and 90°) and 20 trials for each intermediate grating, for a total of 240 trials.

### Task performance and psychophysics analysis

Learning curves across trials were calculated by dividing the number of correct responses (hits + correct rejections) in the preceding 25 trials by 25. Across days the curves were stitched together and smoothed with a Gaussian kernel. If the animals ceased licking for more than 15 trials, the trials were removed from the learning curves, as blocks of inactivity of this size indicate the mouse is distracted or satiated. Discriminative licking was detected by Wilcoxon rank-sum tests on the licking histograms (100 ms bins) generated for each trial (significance for p<0.01) comparing horizontal trials (<45°) with vertical trials (>45°) at each time bin. The first bin with a significant difference was taken as the ‘discrimination time’. Psychometric functions in **Fig. 1D** were taken from 2 days of task performance (480 trials). The criteria for selecting these days was that total performance was above 70% correct across the entire day and there were not too many false alarm trials at the beginning of training (indicating over-thirst) or too many miss trials at the end of training (indicating satiation).

### Cortical lesions and histology

After all mice learned to discriminate horizontal and vertical gratings (n=5 mice) and some learned the full psychometric version of the task (n=2 out of the 5), they were anesthetized (1.5% isoflurane delivered with Somnosuite, Kent Scientific) and placed in a nose clamp. A thermal blanket kept body temperature above 36° C. Ocrygel (TVM Lab) was applied the eyes to keep them from drying out. The location of the C2 barrel had been marked on the skull (A/P: −1.5mm, M/L: 0/3.3mm) from the headpost implantation surgery in these mice, and this mark was used as the center of a 3-4 mm diameter craniotomy. Thermo-coagulation lesions were carried out with a fine tipped cauterizer, making sure not to touch the surface of the brain, but to bring the cauterizer just close enough to blacken the exposed cortical tissue containing the barrel field. The craniotomy was then covered with Kwik-Cast (World precision instruments), and then sealed with dental cement. Sham animals underwent the same surgical procedure except they did not receive thermo-coagulation lesions. After surgery and recuperation (~1 hour in a recovery cage), mice were given 250 μL of water and returned to their home cages. The behavioral testing began again the day after surgery. When behavioral testing was complete, lesioned mice were transcardially perfused with saline followed by a 4% formaldehyde solution in 0.1 M phosphate buffer (PB). Brains were dissected and then post-fixed overnight at 4° C. After washing with phosphate-buffered saline (PBS), brains were cut into 80 μm coronal slices. Slices were mounted and then imaged using a Nikon eclipse 90i microscope (Intensilight, Nikon) and Nikon Pan UW objectives (1x/0.04 W.D 3.2 or 2x/0.06 W.D. 7.5). Slices were then manually aligned with the Paxinos mouse brain atlas and the lesioned areas were tracked along the anterior-posterior axis to make sure the covered the posterior-medial barrel field (fig. S2). Sham mice were used later for electrophysiological recordings during task performance, after which their brains were treated in the same way, except they were sliced tangentially to reveal electrode locations with respect to the barrels (fig. S3). Electrode tracks in these preparations were visible because 1,1’-Dioctadecyl-3,3,3’,3’-Tetramethylindocarbocyanine Perchlorate (DiI) was placed on the shanks before they were inserted into the brain.

### Whisker movement tracking

During some sessions, high speed videos of the whisker interactions with the gratings were filmed with an infrared video camera (Baumer, 500 fps). The frames were grabbed on the same clock as the stimulus presentation to assure synchronization. For each session of whisker videos, a region of interest (ROI) was manually selected around the bases of the whiskers that were in focus. In this ROI, the centroid of the binarized whiskers was computed, and this centroid was then projected onto a line that was perpendicular to the rostral whiskers to give a single coordinate. The velocity of the centroid coordinate across frames was rectified and smoothed to give the whisking envelope. This quantifies the global rostral-caudal movement of all the whiskers. This procedure is graphically displayed in **Fig. 2C**. Normalization to whisking levels in the first 30 frames (first 60 ms of a trial) was sometimes applied to compare across mice with different levels of baseline whisking activity.

### Electrophysiological recordings

On the day of the recording, mice were briefly anesthetized (30 minutes, 1% isoflurane delivered with Somnosuite, Kent Scientific) and the dental cement that was covering the craniotomies from the sham surgery (n=5 sham animals) was removed. In 4 other experiments, fresh craniotomies were drilled following the same protocols described in the lesion section above (except no lesions). After durotomy, the exposed cortical surface was moistened with fresh Ringer’s solution and then covered with Kwik-Cast (World precision instruments), which was secured in place with cyanoacrylate. The mice were then allowed to recover for 2-3 hours in a cage that was placed on a heating blanket. Mice were then placed in the behavioral setup and the Kwik-Cast was carefully removed, making sure not to damage the brain in the process. Multi-electrode silicon probes (A2×32 5mm-25-200-177, Neuronexus) that had been coated with DiI were then slowly lowered into the left hemisphere barrel cortex at about 2 μm per second. Once they reached a depth of 800-1000 μm and sufficient spiking activity was seen across all channels. The preparation then stabilized for 20 minutes before the behavioral protocol was launched, with periodic water rewards given to keep the mice awake and unstressed. In 5 mice, intermediate orientations were rewarded or punished and the number of trials for each orientation followed the protocol detailed in the orientation discrimination training section. In 4 mice, intermediate orientations were given as catch trials, and in these experiments, fewer intermediate orientation trials were given (90 horizontal trials, 90 vertical trials, and 5 catch trials for each of 4 intermediate orientations). Psychometric data was pooled across these 9 mice for the electrophysiological data set. For the behavior alone (**Fig. 1**), all animals followed the same protocols that are described in the orientation discrimination training section.

### Data processing and analysis for electrophysiological recordings

Extracellular signals were acquired at 20kHz with an Intan RHD2000 recording system. The raw data was median filtered to remove common mode noise from all channels and then passed into KiloSort2 for spike detection and clustering. Clusters were manually curated to pick out waveforms with physiological shapes that decay with distance from a primary electrode (electrode with the largest magnitude waveform). The units that passed visual inspection and entered the analysis pipeline were both single units and multi-units depending on the refractory periods found in their autocorrelograms. Data from single and multi-units was pooled for all analyses. Trial-averaged spiking histograms were created by binning spikes in 50 ms bins (**Fig. 3**). Normalized firing rates were computed by dividing by the baseline firing rate, which was taken as the mean firing rate across 500 ms beginning 1 second before trial start.

### Orientation tuning and response detection

Orientation tuning curves were constructed by breaking trials up into 500 ms blocks. For each unit and each 500 ms block, the total number of spikes for a stimulus of a given orientation determined the magnitude of the vector pulling in that direction in a polar coordinate system where all the orientation angles were multiplied by 2. The vector sum of these 10 (or 6) oriented vectors (0, 18, 36, 54, 72, 108, 126, 144, 162, 180° or 0, 36, 72, 108, 144, 180°) was compared to the distribution of vector sums obtained by shuffling the trial labels 200 times. If the actual vector sum was outside of the sphere defined by 95% of the 200 shuffles (p<0.05), then the cell was called orientation tuned in that 500 ms block. False positive rates were thus kept at 5%. Cells were deemed significantly responsive if evoked firing rates were 5 standard deviations above the baseline firing rate.

### Defining the exploratory period and the after-feedback period

Significant differences in licking behavior were assessed by binning the digital lick signal counts into 100 ms bins. Then, the distributions of Go trial licks and NoGo trial licks were compared at each time point relative to trial onset using Wilcoxon rank sum tests, and the first time point in the trials that gave a significant result with p<0.01 is where the mouse was said to have licked discriminatively. For each mouse, the 500 ms before this time point was counted as the early period. The late period was a fixed period after reward or punishment that was chosen to maximize differential licking behavior, after licking had ceased for False alarm trials. This was purposefully chosen to show how a code polluted by drastic differences in licking behavior would present itself.

### Co-firing Analysis

Spikes were binned into 100 ms bins, and the 5×100ms bins leading up to grating halt were used to calculate co-firing. To do this, for each pair of units, the dot product of their activity profiles (5 bins of activity on a given trial) was computed and averaged across trials of a stimulus type (horizontal or vertical). Then the difference was computed between trial types (vertical – horizontal), and this matrix was manually clustered into neurons with different co-firing levels for the respective stimulus classes.

### Support vector machine (SVM) classifiers

Spikes were placed into 100 ms bins to generate population vectors of various types for each trial (**Fig. 5A**). The trials were divided either by grating orientation (**Fig. 5**) or by trial outcome (**Fig. 6**). Binary non-linear SVMs were then trained using the sklearn module in python along with the leave one out protocol in the model selection subdirectory of this module. The non-linear classifiers used a gamma function with an input parameter of 1/n_features (the ‘auto’ option from the sklearn documentation). The population vectors were moved 1 step at a time, and for each time step of 100 ms in the trials, the classifiers were retrained based on the corresponding subspaces of the population vectors that ended at that time step. The performance was the percentage of all trials correctly classified. Each trial was left out only once.

### Paired bootstrap resample test

For small sample sizes (n=5) that are common in challenging experimental conditions such as these, the most appropriate statistical test is non-parametric bootstrap resampling. Wilcoxon and Mann-Whitney tests are often inappropriately used and demand larger sample sizes (n>20). To carry out this test, we resampled 1000 times with replacement from the pool of N (usually 5) mice and permuted the labels of what was being tested (lesion vs. sham, temporal decoders vs. average firing rate decoders, etc). When appropriate, the permutations were done while keeping the measurements paired. If the difference of the mean values obtained was > or < 95% of the shuffled resampled mean differences, then the measurement was deemed significant with p<0.05. Exact p-values are provided as averages of 5 different resamples comprised of 1000 shuffles each.

## General

The authors thank Valérie Ego-Stengel and Isabelle Férézou for giving insightful advice and feedback, Guillaume Hucher for performing the histology, and Aurélie Daret for providing the animal care for the mice.

## Author contributions

ERH built the experimental setup, developed the behavioral task, carried out behavioral experiments and electrophysiological recordings, did analysis, prepared the figures and wrote the manuscript. AR did behavioral experiments, analysis, and edited the manuscript. BB led the project, oversaw the analysis, and wrote the manuscript..

## Data and materials availability

All raw data and code is available upon request.

## Supplementary Materials

**Figure S1.**
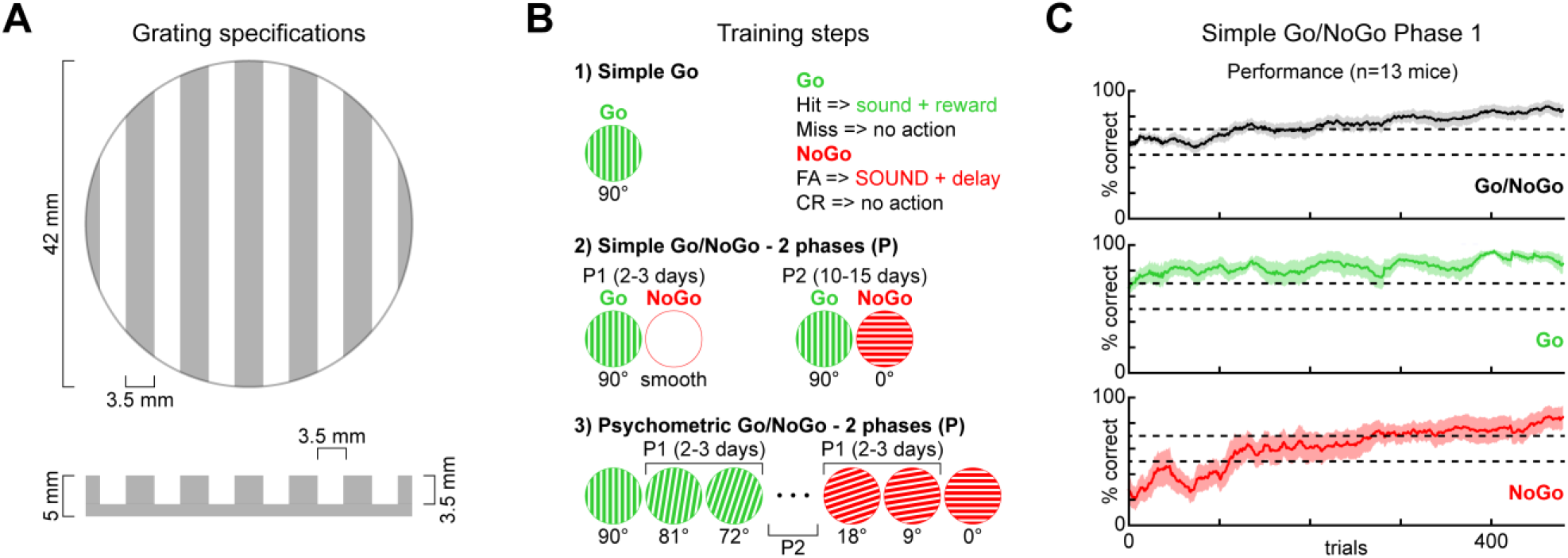
Grating details and behavioral training progression. **A.** A schematic giving the precise dimensions of the gratings. **B.** Task parameters for (1) simple Go detection (2) simple Go/NoGo discrimination with either a smooth surface (Phase 1) or an orthogonal grating (Phase 2) and (3) psychometric Go/NoGo grating orientation discrimination with some intermediate orientations (Phase 1) or all intermediate orientations except 45° (Phase 2). **C.** Mean learning curves for simple Go/NoGo discrimination with a grating and a smooth surface (shaded areas are s.e.m, n=13 animals) for All (black), Go (green), and NoGo trials (red).

**Figure S2.**
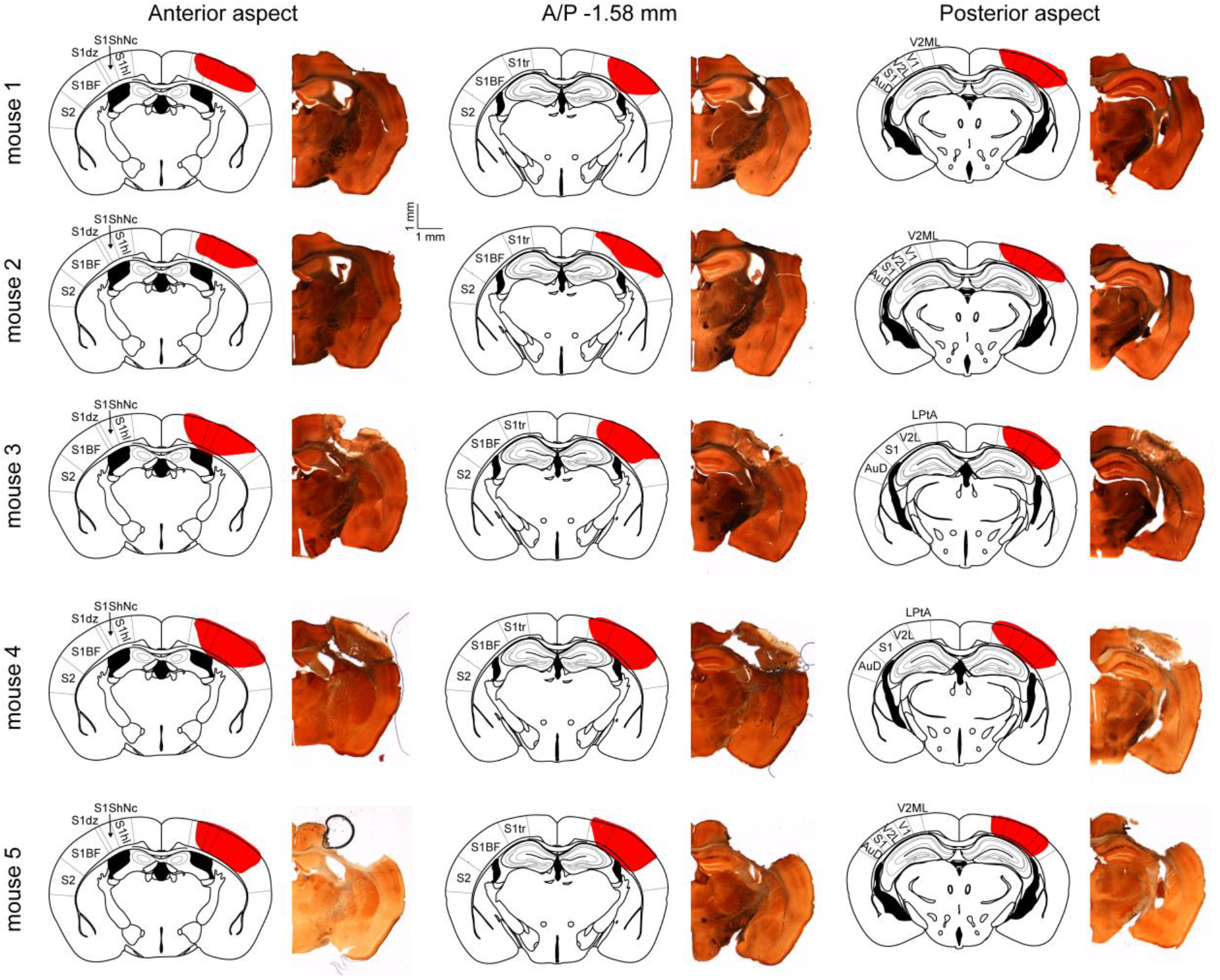
Histology to verify the extent of cortical lesions. Left: Coronal sections from the Paxinos Atlas at the anterior aspect of barrel cortex. They were aligned manually with the histological preparations to the right from each lesioned mouse. The red shading is the lesioned section of the cortex. Center: The same but for the coronal sections that include the coordinates we used to target the center of the posteromedial barrel field. Right: The same but for the posterior aspect of barrel cortex. S1 – primary somatosensory cortex, BF – barrel field, S2 – secondary somatosensory cortex, dz – dysgranular zone, tr – trunk, hl – hindlimb, ShNc – shoulder/neck, AuD – secondary auditory cortex dorsal, V2L – secondary visual cortex lateral, V1 – primary visual cortex, V2ML – secondary visual cortex mediolateral, LPtA – lateral parietal association cortex.

**Figure S3.**
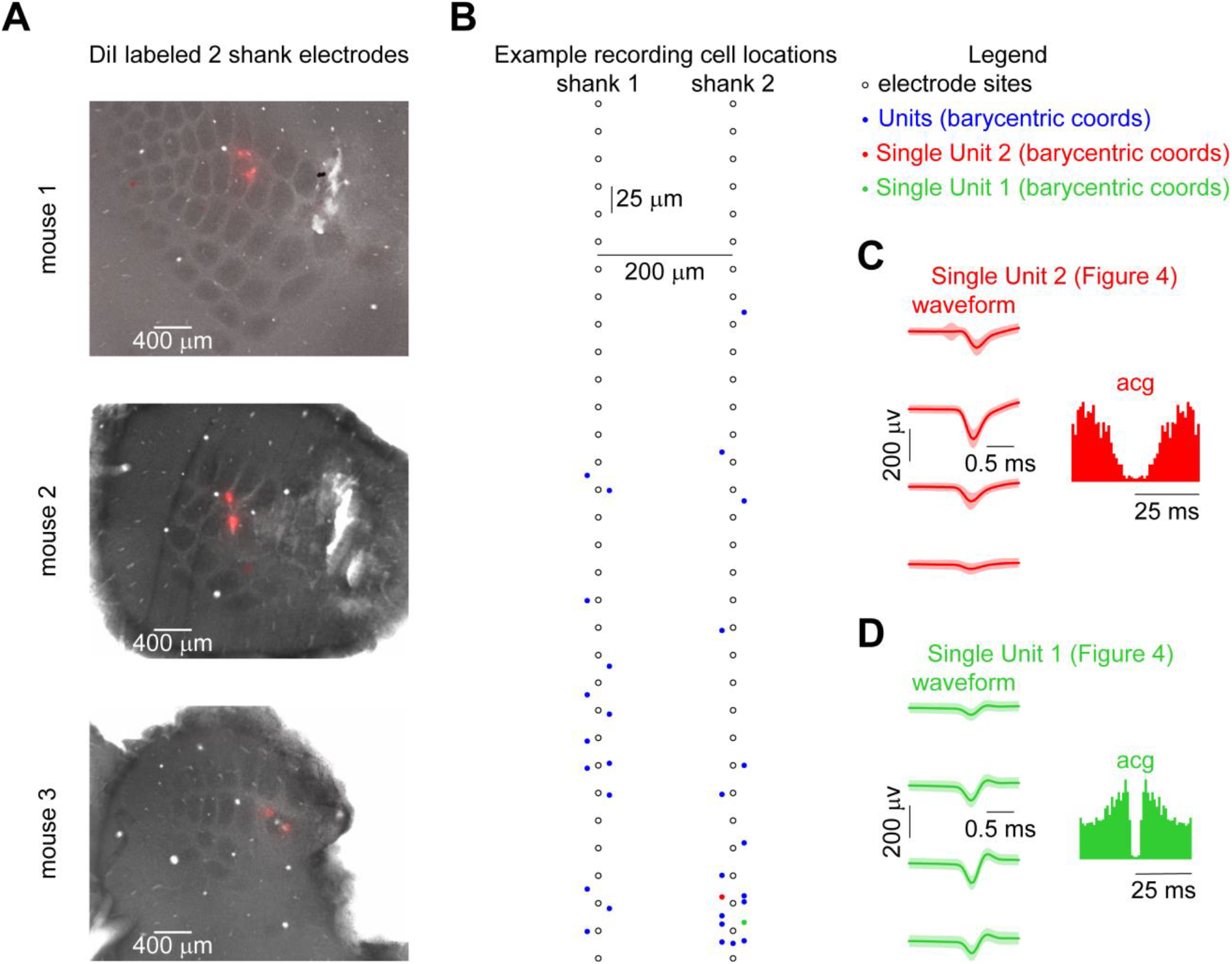
Supplemental information for extracellular recordings. **A.** Electrode placements for 3 example animals showing DiI stained areas within the barrel region of primary somatosensory cortex that was labelled with cytochrome oxidase. **B.** An example recording with the location of every cell mapped onto the electrode sites based on the energy of the waveforms at each electrode position (see Methods). **C-D.** Waveforms and autocorrelograms for the units presented in **Fig. 4**.

**Figure S4.**
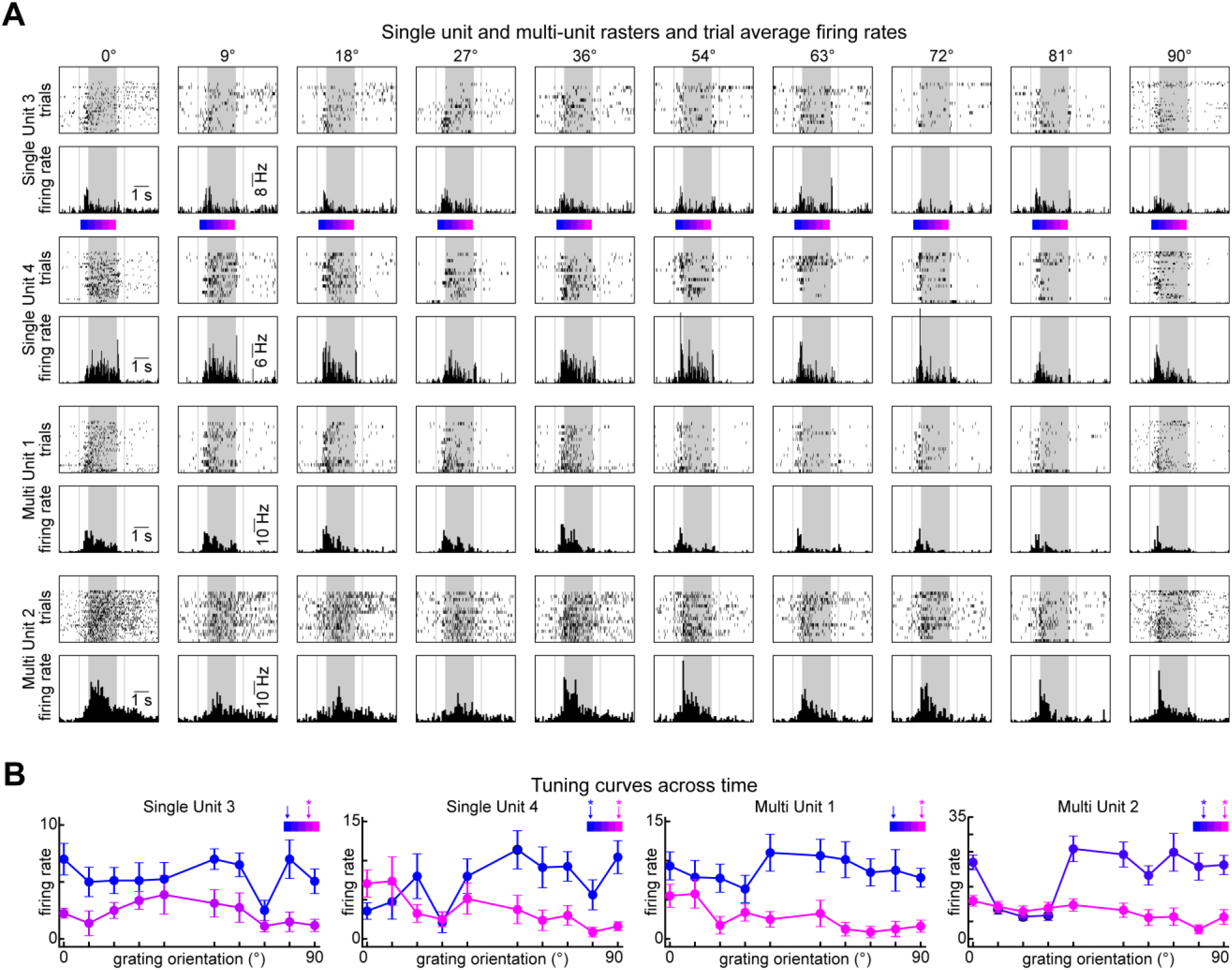
Grating orientation tuning in example single and multi-units. **A.** Single trial rasters and trial-averaged PSTHs from two example single units (top 2) and two example multi-units (bottom 2) during performance of the psychometric Go-NoGo task. **B.** Tuning curves from example units (same units from a) in various time windows color-coded by where the 500 ms time bin in which the curves were computed falls with respect to early vs. late periods. Asterisks indicate significant orientation tuning according the shuffling test (See Methods).

**Figure S5.**
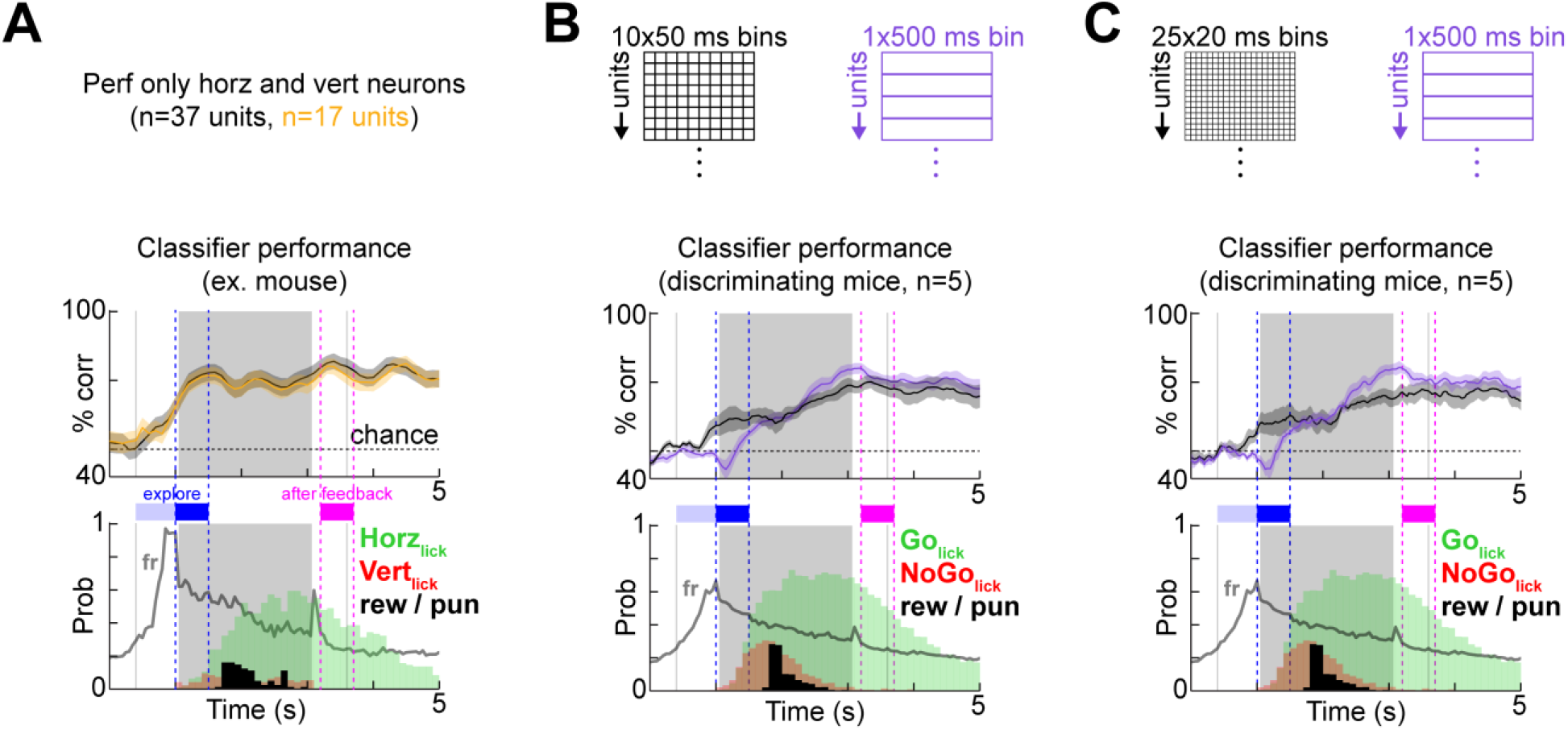
Performance of temporal decoders is robust to changes in bin size and cell dropout. **A.** The classifier performance is the same in an example mouse if it is only trained on a subpopulation of units that have orientation-specific co-firing behavior (with **Fig. 5A-B**). **B.** Two classifier types defined by their bin arrangements: temporal (black, bin size is 50 ms) and average firing rate (purple). Performance, licking behavior and population firing rates are shown for the discriminating (left, n=5) group of mice. **C.** Same as B but for 25 ms bins.

